# A colorimetric comparison of sunless with natural skin tan

**DOI:** 10.1101/2020.05.14.095778

**Authors:** Kinjiro Amano, Kaida Xiao, Sophie Wuerger, Georg Meyer

**Affiliations:** Department of Psychology, University of Liverpool, Liverpool, L69 7ZA, United Kingdom; School of Design, University of Leeds, Leeds, LS2 9JT, United Kingdom

## Abstract

The main ingredient of sunless tanning products is dihydroxyacetone (DHA). DHA reacts with the protein and amino acid composition in the surface layers of the skin, producing melanoidins, which changes the skin colour, imitating natural skin tan caused by melanin. The purpose of this study was to characterise DHA-induced skin colour changes and to test whether we can predict the outcome of DHA application on skin tone changes.

To assess the DHA-induced skin colour shift quantitatively, colorimetric and spectral measurements of the inner forearm were obtained before, four hours and 24 hours after application of a 7.5% concentration DHA gel in the experimental group (*n* = 100). In a control group (*n* = 60), the same measurements were obtained on both the inner forearm (infrequently sun-exposed) and the outer forearm (frequently sun-exposed); the difference between these two areas was defined as the naturally occurring tan. Skin colour shifts caused by DHA tanning and by natural tanning were compared in terms of lightness (*L*\*), redness (*a*\*) and yellowness (*b\**) in the standard CIELAB colour space. Naturalness of the DHA-induced skin tan was evaluated by comparing the trajectory of the chromaticity distribution in (*L*\*, *b*\*) space with that of naturally occurring tan. Twenty-four hours after DHA application, approximately 20% of the skin colour samples became excessively yellow, with chromaticities outside the natural range in (*L*\*, *b*\*) space. A principal component analysis was used to characterise the tanning pathway. Skin colour shifts induced by DHA were predicted by a multiple regression on the chromaticities and the skin properties. The model explained up to 49% of variance in colorimetric components with a median error of less than 2 ΔE. We conclude that the control of both the magnitude and the direction of the colour shift is a critical factor to achieve a natural appearance.

## Introduction

Skin tanning is popular in most western cultures [1], despite the known health risks of UV radiation [2]. One way to mitigate the health risks associated with UV exposure is the use of sunless tanning products. The main ingredient of these artificial tanning products is dihydroxyacetone (DHA) [3], which reacts with the protein and amino acid composition in the surface layers of the skin, keratin and sweat [4]. This reaction leads to the production of melanoidins, a chemical which changes the skin tan to brownish, thereby imitating the effect of the naturally occurring melanin on skin colour [5, 6].

The growing popularity of sunless DHA tanning associated with skin tans being considered as healthy and socially attractive, particularly in the adolescent White population [1, 7–9] necessitates the resulting colour to be close to a “natural” tan (e.g., not too yellow or too orange). Despite of its commercial success, the direction and magnitude of the DHA induced colour change has not been fully characterised. Muizzuddin et al. [10] evaluated skin tonality brought about by DHA with skin tone changes under natural sunlight for Fitzpatrick skin types I-IV (pale-white to light-brown skin) [11]. Based on colorimetric measurements of a natural suntan, a “natural universe of tan” was established in a standard CIELAB colour space based on the measurements of skin tan with 70 White volunteers after the exposure to natural sun light (noon day sun over 72 hours in Long Island, NY in July) and was compared to skin tans changes induced by DHA. Similarly, Chardon et al. [12] determined, on the basis of measurements on 278 White volunteers, a “skin colour volume” defining the chromaticity range of naturally occurring skin tans in a standard CIELAB colour space. The angle in the plane spanned by Lightness (*L*\*) and yellowness (*b*\*) is defined as the individual typology angle (ITA) [12]. The ITA allows a meaningful comparison of skin tone between ethnicities and can be related to the biological mechanisms underlying skin colour [13, 14].

The present study extends previous works [10, 12] (1) by providing a quantitative description of the DHA induced skin tonality changes in terms of colour direction and magnitude, which is then compared to the naturally occurring range of White skin colours, including the ITAs as defined by Chardon et al. [12], and (2) by predicting the magnitude and direction of the colour changes due to DHA from the original skin colour.

To assess whether the DHA-induced skin colour shifts are consistent with naturally occurring tan, colorimetric and spectral measurements were obtained before application of DHA, four hours and 24 hours after DHA application. Our main finding is that, while the trajectory of a *natural* tan is characterised by a decrease in lightness without an increase in yellowness, the direction of skin colour change induced by DHA follows a straight line in the lightness vs. yellowness diagram, leading to an excess of yellowness for a particular lightness level. The result suggests that the control of the colour shift magnitude is a critical factor to achieve a natural appearance by the artificial DHA tanning.

## Methods

### Participants

#### Experimental Group

One hundred staff members and students at the University of Liverpool (female: *N* = 83, range = 20-44 years, median = 23; male: *N* = 17; range = 18-44 years, median = 27) participated in the experiment. Ninety-four out of 100 were White (female 78, male 16) and six were non-White (female five, a male) consisted of one Mixed, five Asian/Asian British (including two individuals with darker skin). More than a half of the participants (66 out of 100) reported having used self-tanning products before, but none of them had used the tanning products within the five weeks prior to taking part in this study. Exclusion criteria included pregnancy; evidence of acute or chronic disease, and dermatological problems. Participants’ ethnicity was not used as an inclusion/exclusion criterion for the recruitment but age range was limited between 18 and 45 years. There were no restrictions on physical activities and types of diet during the participation, but participants were advised not to scrub the test area hard while showering and not to apply any skincare products on the area. All participants gave informed consent. Informed consent was obtained as part of an online demographic questionnaire and stored in a secure server. The study was approved by the University of Liverpool Ethics Committee (IHPS 4746), consistent with the Helsinki Declaration. The measurements were carried out from March to May 2019 at the University of Liverpool.

#### Control Group

A second data set of skin colour measurements was obtained from a separate cohort [15], as part of the Leeds-Liverpool skin colour database (CIE Technical Committee TC 1-92 Skin Colour Database [15, 16]) which includes measurements of ten body locations: six facial (forehead, cheekbone, cheek, nose tip, chin, and neck) and four arm locations (the back of the hand, inner forearm, outer forearm, and fingertip). For the purpose of this study, we only made use of the inner and outer forearm measurements of 60 White (female 46, male 14; range = 18-39 years). This cohort of the 60 White is labelled as a “control group” in the following sections.

### Materials

A generic self-tan bronzing water-based gel with a 7.5% DHA concentration as active ingredient was used. The gel contained no penetration enhancers or other materials that affect reactivity of the DHA. All ingredients in the product are in line with EU Cosmetic Regulations and are as follows: Aqua (Water), Dihydroxyacetone, Glycereth-26, Hydroxyethylcellulose, Phenoxyethanol, Benzyl Alcohol, Sodium Metabisulfite.

### Skin colour measurements

Skin measurements for each participant in the experimental group were taken on three occasions over two consecutive days: before the DHA gel application, as a baseline, at four and 24 hours after the application. In the following sections data of the baseline measurement are labelled as “0h”, four hours after the application as “4h”, and 24hours after the application as “24h”.

All measurements were taken on the inner arm (relatively infrequent exposure to sunlight). On each test area three points were randomly chosen, while avoiding skin spots, freckles, or moles in the area. Because the difference between the right and left arms are very small, results and analysis are based on the average values over all six measurements per participant. The measurements were taken with a spectrophotometer (model CM-700d, Konica-Minolta, Tokyo, Japan), with an 8 mm aperture and low pressure mask, under illuminant D65 with the CIE 2 degree standard observer; spectral range from 400nm to 700nm in 10nm step. The aperture of the photometer was placed on the skin surface to take the measurements. The low pressure mask enabled stable measurements without applying excessive pressure on the surface [17]. Within this project, the experimental group was measured with the exact same parameter settings and the same protocol as the previously measured control group [15–18].

All colorimetric skin measurements will be presented in a standard CIELAB colour space with three axes: lightness (*L*\*), redness (*a*\*) and yellowness (*b*\*), where chroma is defined as the length in (*a*\*, *b*\*) space [19, 20]:

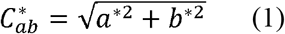

To quantify the overall colour shift, the CIELAB colour difference metric Δ*E* [19, 20] from the baseline 0h to 4h and 24h were calculated. Colour difference Δ*E* represents the perceptual difference between the two colour samples. It is defined as the Euclidian distance between two colours in a perceptually uniform colour space (i.e., CIELAB colour space).

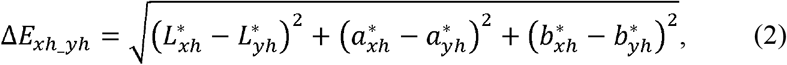

where *x*_h_ corresponds to 4h or 24h and *y*_h_ to the baseline 0h. Δ*E* at unity corresponds to a just-noticeable colour difference in a controlled viewing environment.

#### Skin colour classification

We used Chardon et al.’s [12] index to classify the skin colours in CIELAB space. The Individual Typology Angle, ITA [12] (see also, [13, 14]) is defined as an angle in the (*L*\*, *b*\*) diagram, with the origin at *L*\* = 50 (Fig 1A):

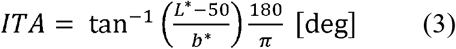

**Fig 1.**
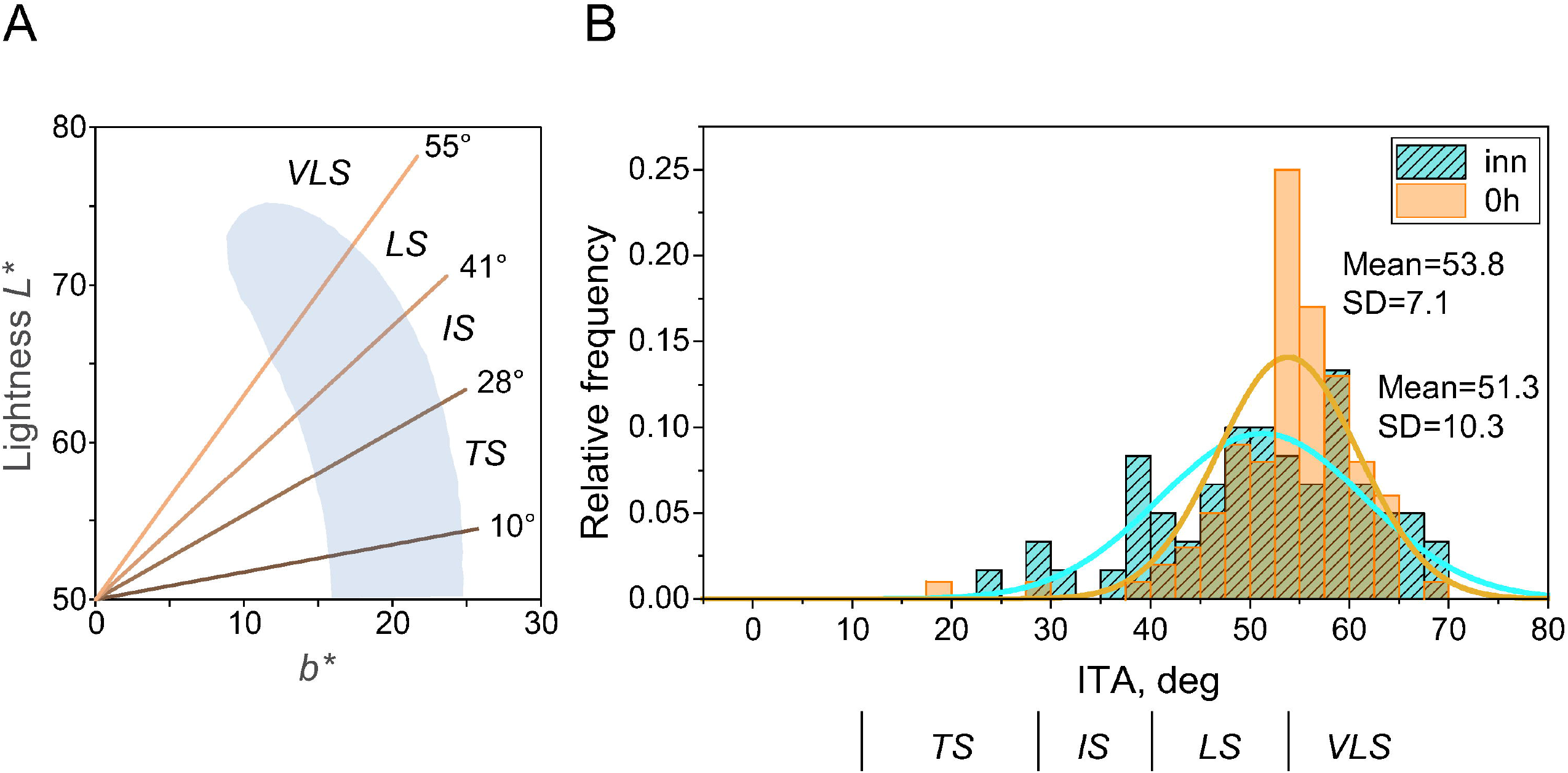
(A) Skin colour volume defined in lightness-yellowness space (*L*\*, *b*\*). The shaded area depicts the Chardon’s skin colour volume of a large White sample [12]. The Individual Typology Angle (ITA) is defined in Equation (3). Chardon et al.’s [12] skin colour categories “very light” (VLS), “light” (LS), “intermediate” (IS) and “tan” (TS), VLS > 55° > LS > 41° > IS > 28° > TS > 10° are indicated (see also, [13]). (B) Histogram of relative frequency of ITA for the experimental group (inner forearm at 0h, *n* = 100) and the control group (inner forearm, *n* = 60). Chardon et al.’s [12] skin colour categories are indicated underneath of the abscissa.

Skin colour is classified by ITA as “Very Light Skin” (VLS), “Light” (LS), “Intermediate” (IS) and “Tan” (TS), using the following categories: VLS > 55° > LS > 41° > IS > 28° > TS > 10°. The light grey area (“skin colour volume”, reproduced from Fig 6, ibid) shows the envelope of all natural skin colours found in the White population, ranging from ‘very light’ pale skin (at the top of the area) to ‘brown’ White skin (the bottom), derived from 278 White volunteers [12]. Naturally occurring lightness levels (*L*\*) range from 50 (brown skin) to 75 (light pale skin), whereas skin yellowness (*b*\*) ranges from 10 to 25 (see also, [13]). Crucially, the maximally occurring yellowness (*b*\* = 25) is reached at a lightness level of *L*\* = 50.

Fig 1B shows the ITA histogram for the experimental group at 0h (‘0h’) and the inner arm of the control group (‘inn’, Fig 1B). These distributions are overlapping with similar means (control group: 51.3°; SD = 10.3; experimental group: 53.8°; SD = 7.1) which are not statistically different, as confirmed by a two-sample *t*-test (*t*(158) = 1.87, *p* = 0.063) and therefore allows direct comparisons between these groups (details in the Results).

### Skin moisture and pH level measurements

A number of factors, such as the DHA concentration, additive ingredients, and environmental attributes have been shown to affect the colouring reaction [21]. Since the pH level of the skin surface affects the stability of DHA [4], it is possible that the pH level modulates the effect of DHA reaction. We therefore monitored the skin moisture and pH level along with colorimetric and spectral measurements. Skin moisture content and pH level were measured using a Corneometer and pH probe, respectively, which are integrated on the CK multi probe system (model CM5, Courage + Khazaka Electronic GmbH, Germany). The measurements of the skin properties, skin moisture and pH were taken at five points across each of the test areas at 0h, 4h and 24h. The average of the ten measurements (for both arms) was used in the subsequent analysis.

### Procedure

#### Experimental group

Prior to skin measurements, the participants answered a questionnaire including her or his attitudes towards self-tanning as well as basic demographic. At 0h, test areas on inner forearms on both left and right inner forearms were identified using a square shaped mask of 5 cm × 5 cm. Colorimetric and spectral measurements were taken at three points within each test area then averaged. Skin moisture and pH level were taken at five points within each test area then averaged. Then one ml of DHA gel was applied uniformly on each test area. The time for the gel to be absorbed was approximately 10 minutes. Participants returned to the laboratory four hours and 24 hours after the DHA application for repeated measurements.

#### Control group

The measurement procedure is described in detail elsewhere [15]: the same instrument and identical instrument settings and protocols were used.

## Results

### Colorimetric shifts induced by sunless tanning (DHA)

Colorimetric data of all 100 participants (experimental group) are represented in CIELAB space in Figs 2A. Different colours represent measurements at 0h, 4h, and 24h, respectively; ellipses represent 95% confidence intervals. The skin colour change from 0h to 4h to 24 hours is shown in the yellowness vs. redness diagram (*b*\*, *a*\*) and in the lightness versus chroma diagram (*L*\*, 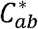) (Fig 2A). Skin colour shifts towards reddish and yellowish, roughly at a rate of two to one in yellowness to redness, in line with the overall direction of variation in the measured sample (the first and second principal components on each cluster of 0h, 4h, and 24h are shown as solid lines). Plotting chromaticities in the (*L*\*, 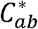) diagram, reveals that the entire distribution shifts downwards showing a negative correlation between lightness and chroma. The area of ellipses in (*a*\*, *b*\*) are 41.7, 36.7, and 33.0 and in (*L*\*, 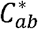) are 77.4, 65.7, and 64.2 for 0h, 4h, and 24h, respectively, demonstrating that the skin colour distributions become tighter with progressive sunless tanning. An example of colour shift on inner forearm by DHA in four and 24 hours are given in Fig 1B.

**Fig 2.**
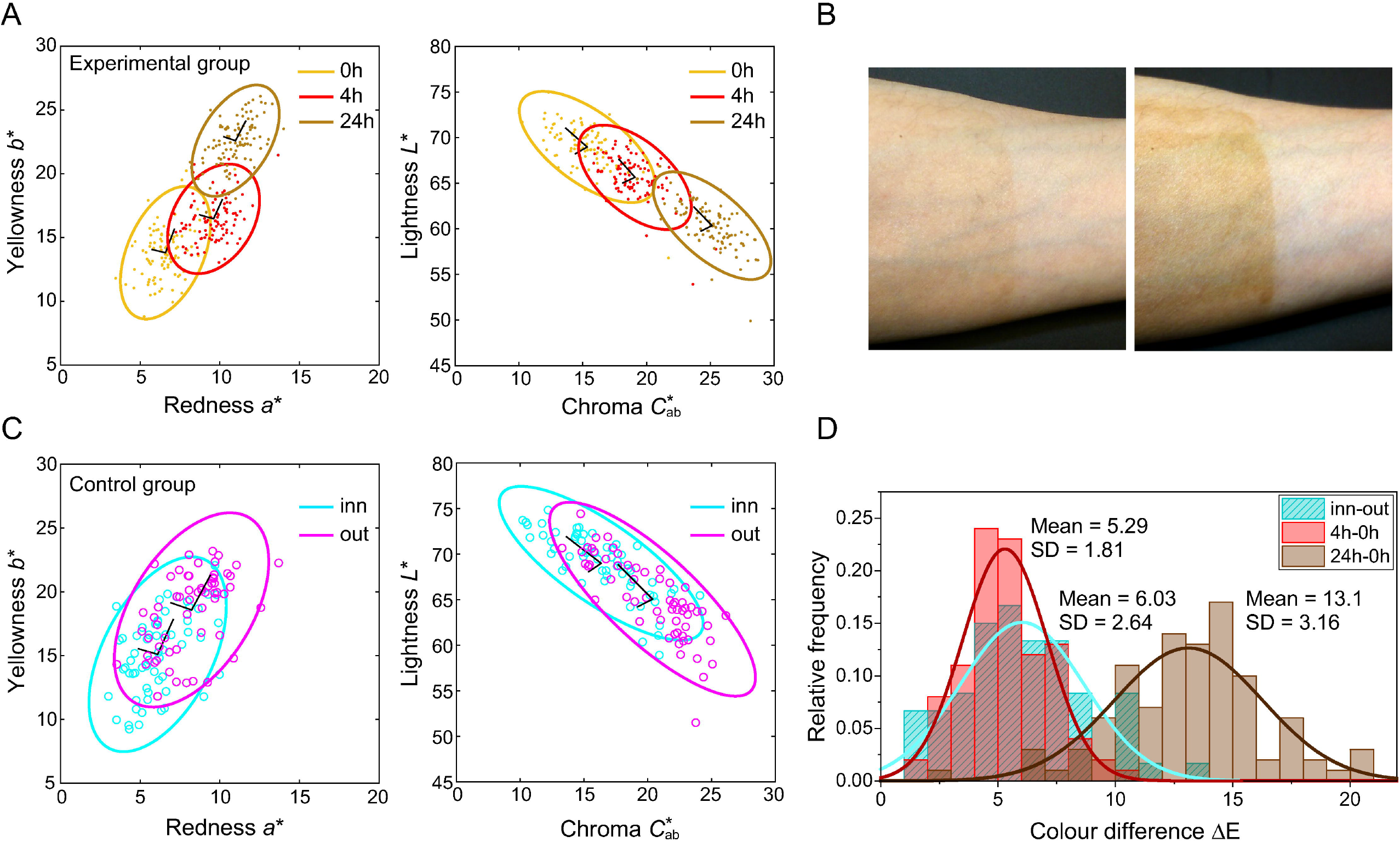
Colorimetric properties of the DHA tanning. (A) Experimental Group. Skin colour distributions in the yellowness vs. redness (*b*\*, *a*\*) and lightness vs. chroma (*L*\*, 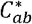) diagrams (experimental group). Each point represents an individual participant. Different coloured symbols indicate the skin colour at 0h (baseline), 4h, and 24h, respectively. Ellipses represent 95% confidence interval of each distribution. The first and second principal components are shown as solid lines; (B). Example of a skin colour change due to DHA at 4h and 24h on inner forearm. (C) Control Group. Skin colour distributions of the inner and outer forearm of the control group. (D) Comparison of the Experimental and the Control Group. Histogram of skin colour differences (CIELAB ΔF) between the baseline (0h) and the DHA tanning after 4h and after 24h compared to the difference between the inner and outer arm (inn-out) of the control group. Solid curves are Gaussian fits.

### Colorimetric comparisons with natural tanning

The colorimetric skin values of the control group are shown in Fig 2C. The comparison of the inner forearm (infrequently exposed to sunlight) with the outer forearm (frequently exposed to sunlight) provides a measure for naturally occurring skin-tan. The distribution of the skin tans of the inner forearm of the control group completely overlap the baseline measurements of our experimental group (0h); the overlap between the skin tan changes of the outer forearm (control group) with the DHA distribution after 4h, is less but still substantial (Fig 2A with Fig 2C). In contrast the DHA-induced skin colours after 24 hours are darker and of higher chroma compared to naturally tanned outer forearm. The histogram of the skin colour differences Δ*E* between the outer and the inner forearm is shown in Fig 2D (‘inn-out’) together with the distribution for 4h-0h and 24h-0h changes. The distribution of Δ*E_out–inn_*(mean = 6.03, SD = 2.64) has a similar mean to the distribution Δ*E*_4*h*–0*h*_(mean = 5.29, SD = 1.81), consistent with the substantial overlap between these two distributions. The overall colour shift induced by DHA 24 hours application (‘24h-0h’) is about 13ΔE, which is very substantial and clearly visible (cf. Fig 2B).

### Spectral comparison between sunless and natural tans

Surface spectral reflectances in the visible wavelength range (400nm-700nm) were obtained at the same time as the colorimetric measurements. The average spectra (six measurements for each of the 100 participants) are shown in Fig 3 A for each of the three measurements. DHA reduces the amount of light reflected across the entire wavelength range, but the effect is stronger in the short-wavelength range. At the shortest wavelength, 400nm, the reflectance ratio of 4h to 0h is approximately 0.8, and reduces to about 0.4 after 24 hours (Fig 3B). This reduction in the reflection of blue-ish (short-wavelength) light is responsible for the skin to appear more yellow-ish.

**Fig 3.**
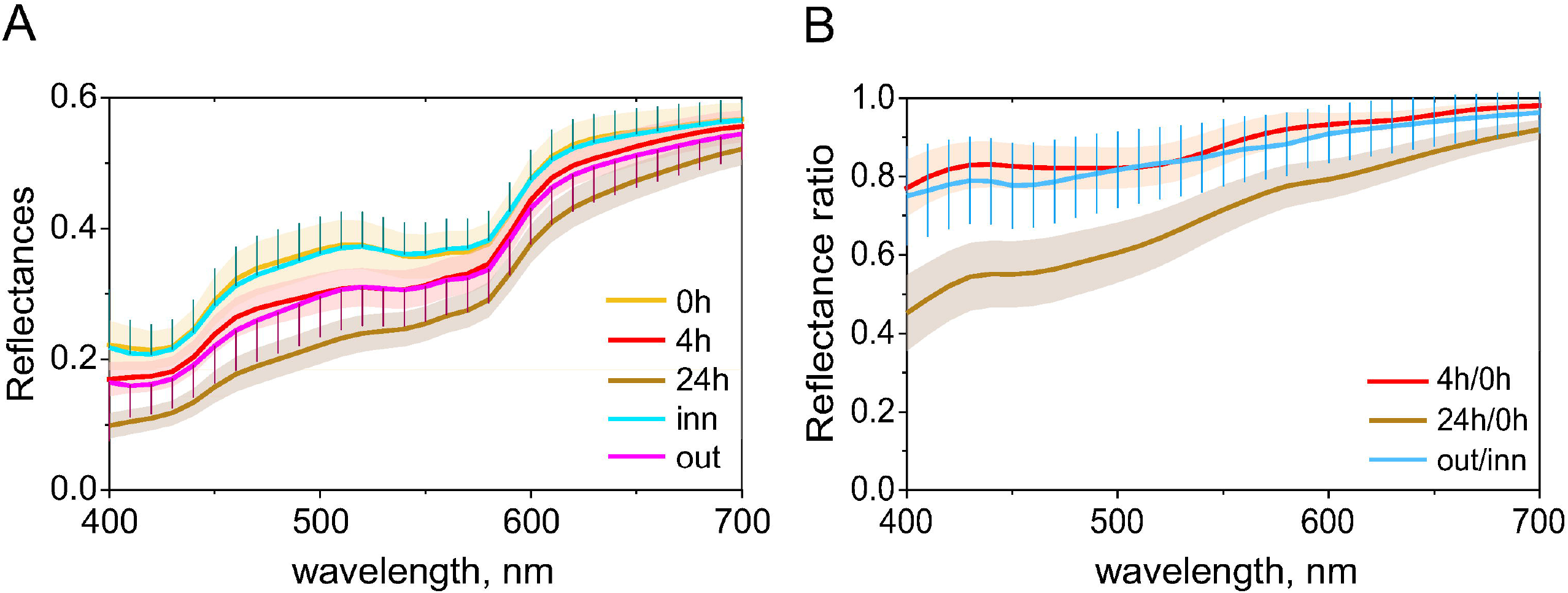
Spectral comparisons of the DHA and the control group. (A) Mean spectral reflectances for the experimental group: baseline (0h), after 4h and 24 hours of DHA application; also the spectra of the inner and outer forearm of the control group. Standard deviations of the two groups; to enhance visibility coloured shade is used for the experimental group while error bars are used for the control group. (B) Reflectance ratios: 0h to 4h, 0h to 24h (experimental group) and inner to outer forearm (control group). Standard deviation is shown with colour shade for the experimental group and error bars for the control group.

The spectral reflectances of the inner and outer forearm of our control group are indicated with ‘inn’ and ‘out’ in Fig 3A. There is a very good agreement between the spectra of the inner forearm of the experimental group at ‘0h’ and the inner forearm of the control group (‘inn’). Four hours after application of DHA, the spectra of the experimental group (‘4h’) are similar to the outer forearm of the control group (‘out’), which is also indicated by the reflectance ratio (Fig 3B): the relative spectral changes after four hours of DHA (4h/0h) are similar to the relative changes seen between the inner forearm (infrequently sun-exposed) and the outer forearm (frequently sun-exposed) of the control group. This is consistent with the colorimetric data showing a substantial overlap in the distribution of the ‘4h’ experimental group and the ‘out’ (outer forearm) of control group (Fig 2A vs. Fig 2C). The similarity of spectral profiles between pairs of spectral samples can be evaluated by RMSE (root mean square error), as it provides better evaluation than the other metrics [22], such as SSV (spectral similarity value) [18, 23]. RMSE is defined as Eq (S1) (Supporting Information). Smaller RMSE indicates that the two spectra are similar.

The entries in the second column in Table 1 show the RMSE calculated from individual participants: thus, RMSE for corresponding participants across the conditions, then averaged over the participants. The entries in the third column were calculated from mean spectral reflectances of each condition (0h, 4h, 24h, *n* = 100; inner and outer forearm, *n* = 60). RMSE between 0h and inn (‘0h-inn’) is very small, confirming the similarity between the two spectra. RMSE between 0h and 24h (0h-24h) was larger than between 0h and 4h (0h-4h), almost the same level as RMSE between inner and outer forearm (inn-out). RMSEs based on the individual participants (the second column) are reasonably close to those based on the mean spectral reflectance (the third column). The same trends were also found with the analysis with SSV (S1 Table, Supporting Information). The RMSE values are consistent with the ITA analysis and show that the spectral differences are smallest between the inner arm of the control group and the experimental group at 0h. Similarly, the experimental spectra at 4h are almost identical to the outer forearm spectra of the control group. Standard deviations (SDs) over participants across spectra (400nm-700nm) became smaller along with the DHA tanning developed (as time progressed from 0h to 4h and 24h) as the time progressed from 0h to 4h and 24h (see S2 Table, Supporting Information).

**Table 1.** RMSE between pairs of spectral reflectances

|  | RMSE |  |
| --- | --- | --- |
|  | individual spectra (SD) | mean spectra |
| 0h-4h | 0.044 (0.016) | 0.044 |
| 0h-24h | 0.108 (0.028) | 0.108 |
| inn-out | 0.052 (0.026) | 0.050 |
| 0h-inn |  | 0.005 |
| 0h-out |  | 0.053 |

### Comparison with the skin colour volume of Chardon et al (1991)

In addition to comparisons with our control group, we now analyse the DHA-induced skin tone changes using the skin colour volume derived by Chardon et al. [12] (details in Fig 1A). (The skin colour volume and ITAs across different ethnicities have been reported in [13]).

Overlaying the DHA skin colours for 0h, 4h, and 24h on the skin colour volume (natural skin gamut, shaded area in Fig 1A) shows that over 20% of the DHA-induced skin colours are outside of the natural gamut after 24h due to excessive yellowness (Fig 4A, upper panel). The DHA-induced colour changes are best described by a straight line in (*L*\*, *b*\*) diagram, whereas the natural tanning pathway follows a curved shape. The skin colours of our control group, as expected, are within the natural skin gamut (Fig 4A, lower panel) and the orientation of the ellipses (inner and outer forearm) are aligned with the ‘banana shape’ of the natural skin gamut instead of following a straight trajectory in the (*L*\*, *b*\*) diagram.

**Fig 4.**
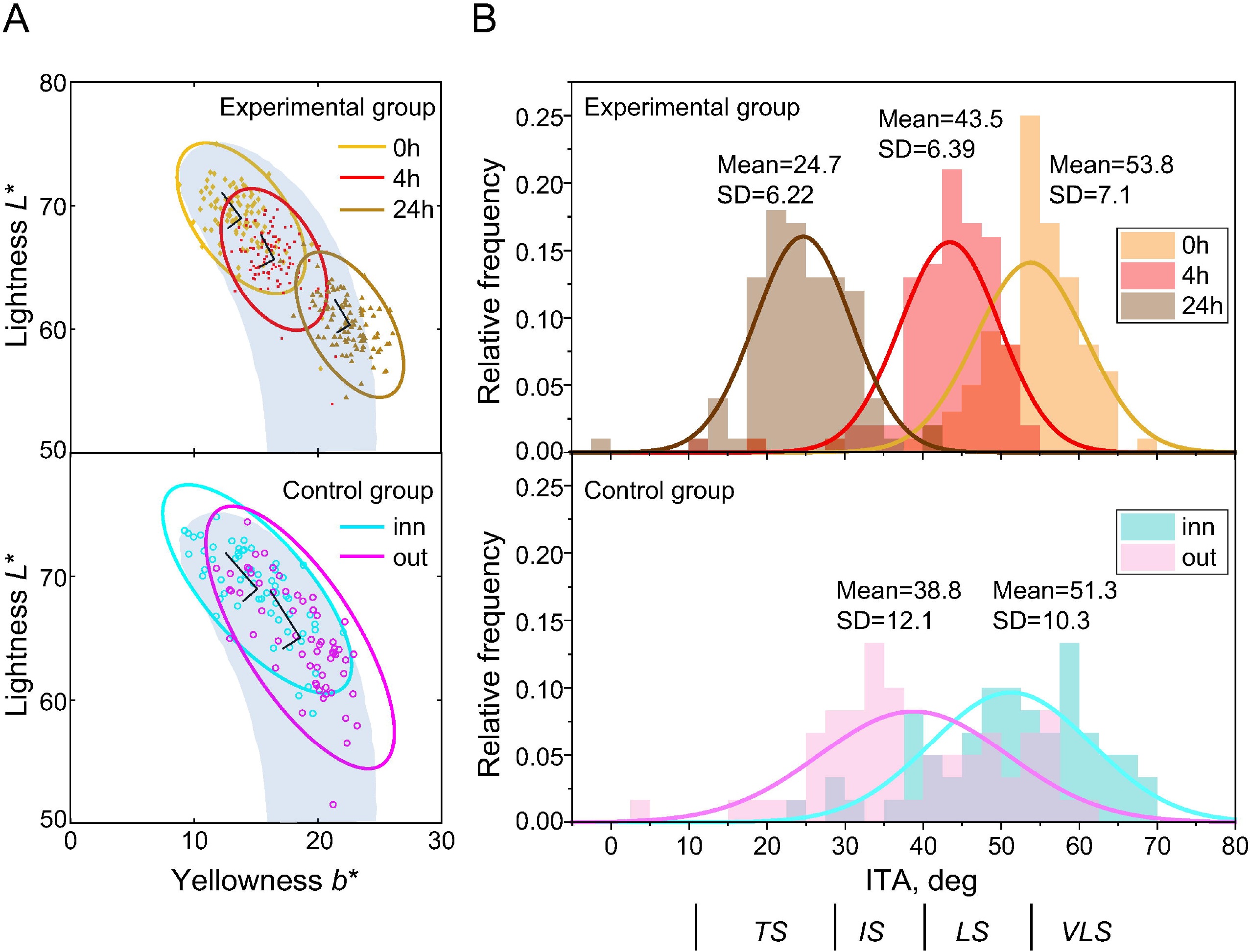
Chromaticity distributions in CIELAB (*L*\*, *b*\*) and ITA distributions. Upper panel show the chromaticity and ITA distributions at 0h (baseline), 4h and 24h (DHA tanning) of the experimental group, whereas lower panel shows those of inner and outer forearm of the control group. (A) Chromaticity distribution in (*L*\*, *b*\*) diagram. Shaded area represents the naturally occurring skin colour gamut, “skin colour volume” of Chardon et al.[12]. (B) Histogram of the ITA (Individual Typology Angle).

The Individual Typology Angle (ITA – see Eq 3 and Fig 1) represents the skin colour in the (*L*\*, *b*\*) diagram. The range of the ITA at 4h (mean = 43.5; Fig 4B) overlap substantially with the ITA of the outer forearm measurements of our control group (‘out’; mean = 38.8), whereas the distribution of the ITA at 24h (mean = 24.7) is shifted towards ‘tanned skin’ (< 28°; lower area of the skin colour volume, see also, Figs 1A and 4A) and largely outside the range of naturally occurring skin tans of our control group (‘out’). The DHA-induced shifts (0h to 4h, 4h to 24h) were significant, as confirmed with a one-way ANOVA and subsequent *t*-tests (one-way ANOVA comparing 0h, 4h, 24h: *F*(2, 297) = 505.4, *p* < 0.05; T-tests: 0h vs 4h: *t*(198) = 10.9, *p* < 0.05; 4h vs 24h: *t*(198) = 30.9, *p* < 0.05).

### Predicting the DHA-induced skin colour changes

We are interested in how each of colorimetric component, (*L*\*, *a*\*, *b*\*), was influenced by the DHA reaction. Therefore we analyse the vector representation (Δ*L*\*, Δ*a*\*, Δ*b*\*) in this section. The shift in skin colour (Δ*E*\*, Δ*a*\*, Δ*b*\*) from 0h to 24h is plotted in Fig. 5A as a function of the baseline colour (*L*\*, *a*\*, *b*\*). For lightness, we find a decrease compared to the baseline lightness (*L*\* at 0h), whereas redness *a*\* and yellowness *b*\* increase after DHA application (cf. Fig 2A). The absolute magnitude of the skin colour shift induced by DHA is not constant, but depends on the initial skin colour, as evidenced by the highly significant negative correlations between the colour shifts and the base colour for each dimension (for *L*\*: Pearson correlation coefficient *r* = −0.47; *a*\*: *r* = −0.51; *b*\*: *r* = −0.67; *p* < 0.001 for all *L*\*, *a*\*, and *b*\*) (Fig 5A).

**Fig 5.**
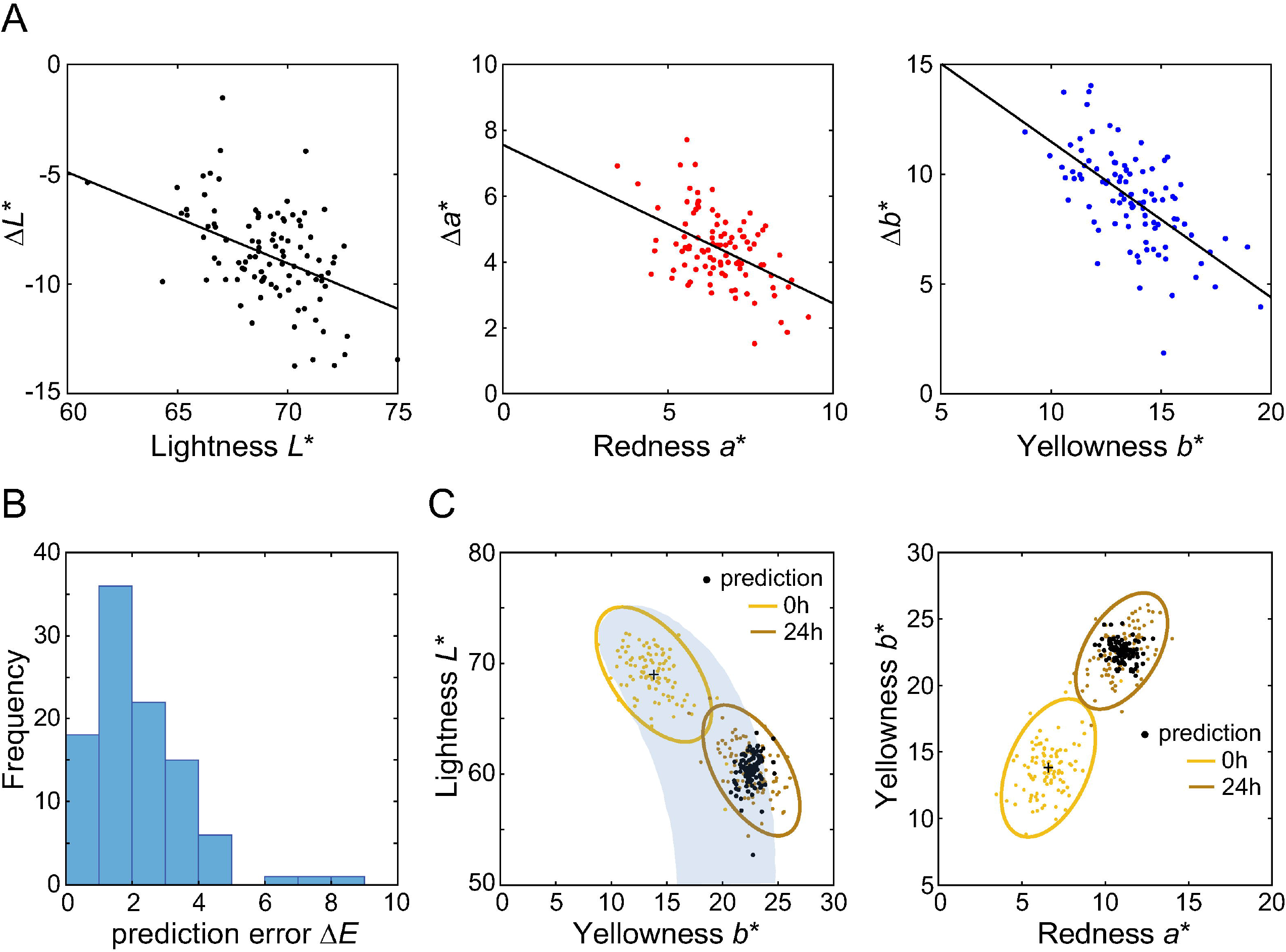
Predicting the DHA-induced colour shifts. (A) The DHA induced changes (24h-0h) are plotted as a function of the baseline colour (0h) for all three colour dimensions. Straight lines indicate linear regression (see text). (B) Histogram of errors defines as the difference between the predicted by a linear model and observed skin colour changes. (C) Predicted skin colour shifts from 0h are indicated as black dots in the (*L*\*, *b*\*) and the (*b*\*, *a*\*) diagram. The shaded area corresponds to the skin colour volume of Chardon et al.[12].

To predict the final DHA-induced skin colour changes (*L*\*, *a*\*, *b*\* in 24h), a stepwise multiple regression (*stepwiselm* function in MatLab2018a; MathWorks Inc, Natick, MA) was performed, with colour change (Δ*L*\*, Δ*a*\*, Δ*b*\*) as dependent variables and the baseline skin colour (*L*\*, *a*\*, *b*\*), skin moisture, and skin pH, and all pairwise interactions (between variables, e.g., between *L*\* and *a*\*, represented by *L*\*. *a*\* in Table 2) as predictors. Stepwise regression adds or removes predictors from the model based on the criterion *p*-value (set to 0.051 here). Table 2 shows the predictors with regression coefficients larger than zero (*p* < 0.01 or *p* < 0.05) in each column and dependent variables in each row. Overall, all three regressions were significant at *p* < 0.001. Neither skin moisture nor the skin pH had any predictive power for the skin colour shifts. Table 2 contains the standardised regression coefficients, the adjusted *R^2^* and degrees of freedom. For the lightness shift (Δ*L*\*), 22% of the variation is explained by *L*\* and no other predictor contributes significantly. Similarly, 27% of the variation in the redness shift (Δ*a*\*) is predicted by *a*\* and *b*\*; shifts along the yellowness dimension (Δ*b*\*) are predicted by *L*\*, *b*\* and their interaction, explaining 49% of the variance.

**Table 2.** Multiple regression analysis for skin colour change

| Dependent variable | Regression Coefficients | | | | Adjusted $R^2$ (df) |
| --- | --- | --- | --- | --- | --- |
| | $L^*$ | $a^*$ | $b^*$ | $L^*.b^*$ | |
| $\Delta L^*$ | -0.47** | - | - | - | 0.22 (98)** |
| $\Delta a^*$ | - | -0.42** | -0.20* | - | 0.27 (97)** |
| $\Delta b^*$ | 0.31** | - | - | -0.09* | 0.49 (96)** |
|  |  |  | 0.58** |  |  |

\*indicates coefficients with $p < 0.05$
\*\*indicates coefficients with $p < 0.01$

For colour application research, it is often useful to understand the prediction error in perceptual units (the CIELAB colour difference Δ*E*, see Eq (2)) instead of variance explained (*R*^2^). Thus, the goodness of the prediction was estimated by calculating Δ*E* from the errors between the predicted and observed values in each of Δ*L*\*, Δ*a*\*, and Δ*b*\*. The smaller the error (Δ*E*) is, the better the prediction would be. The histogram of the prediction error Δ*E* using the linear model (see Table 2) is shown in Fig 5B; the median error is 1.87 where 96% of the error was below 5Δ*E*. For comparison, if we include all 15 predictors (*L*\*, *a*\*, *b*\*, skin moisture, and skin pH as well as all interactions between the all predictors) the median Δ*E* is reduced by about 10% to 1.58.

If we assume that each individual is affected in the same way (using a constant model, instead of a linear model) the median Δ*E* increases to 2.18, almost 20% larger than the linear model (Table 2). The best fitting constant shifts are (−8.6, 4.4, 8.8) in (*L*\*, *a*\*, *b*\*). The predicted skin colours (using the linear model, cf. Table 2) are shown as black dots in Fig 6C with the (*L*\*, *b*\*) and the (*b*\*, *a*\*) diagrams. It is clear that the colour shift induced by DHA is not constant, but depends strongly on the original skin colour as evidenced by the regression analysis (Table 2).

**Fig 6.**
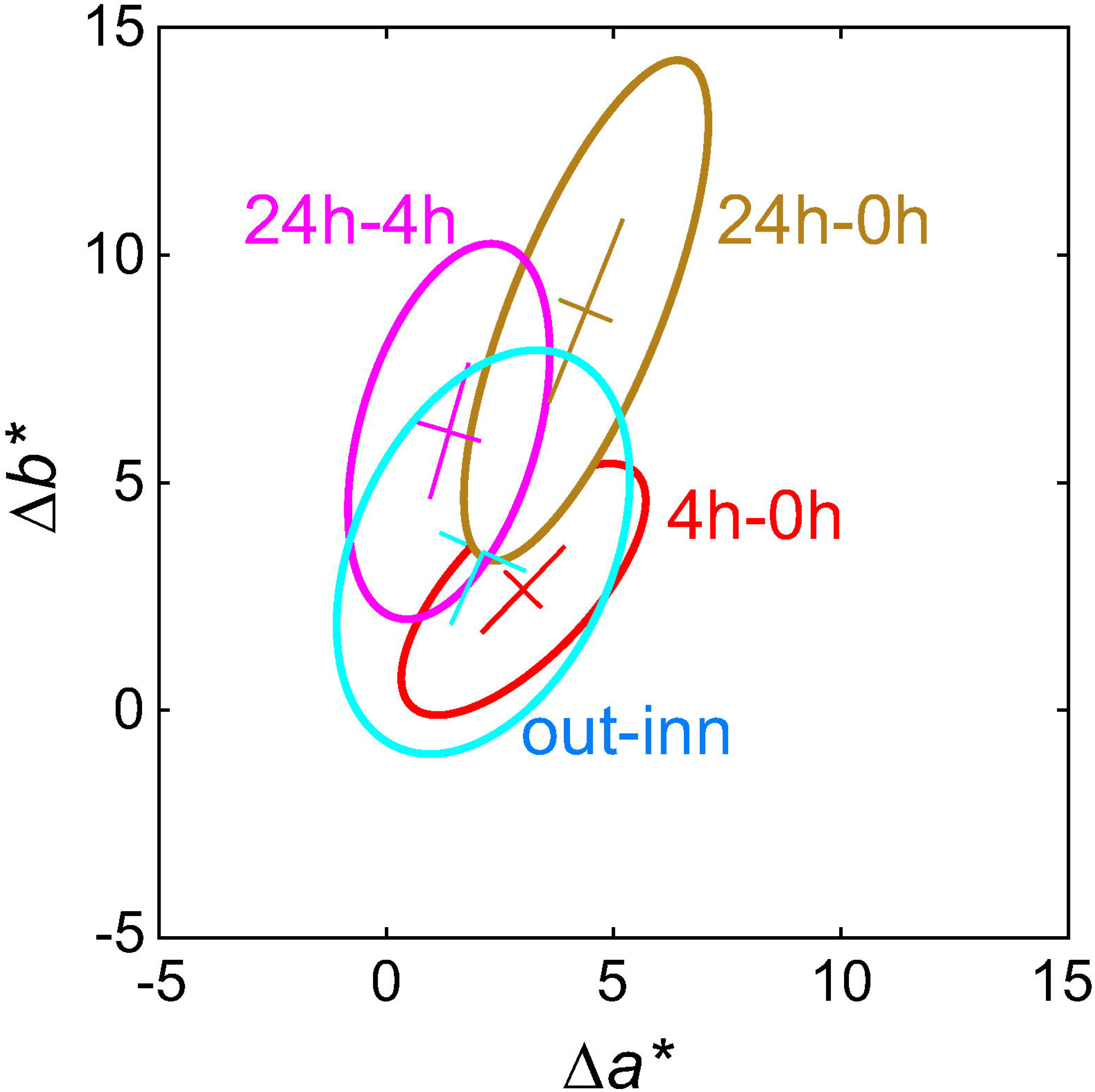
Principal component analysis on chromaticity differences (Δ*a*\*, Δ*b*\*). The shift ‘24h-0h’ is simply the vector summation of ‘4h-0h’ and ‘24h-4h’.

## Discussion

The purpose of the study was to characterise the tanning pathway of a sunless DHA-induced skin tan with the aim to understand when a sunless tan looks unnatural, and whether we can predict the outcome of DHA application on skin tone changes.

### Skin colour changes induced by DHA

Our main finding is that the direction of the skin colour change induced by DHA follows a straight line in the lightness vs. yellowness (*L*\*, *b*\*) diagram, leading to an excess of yellowness for low lightness levels (Fig 4A), while the trajectory of a natural tan is characterised by a curvilinear relationship between lightness and yellowness, with a maximum yellowness at a lightness of 50.

The skin colour volume for White is depicted as the ‘banana shape’ in the natural skin gamut proposed by Chardon et al. [12] (Fig 4A): approximately 20% of the DHA-induced skin colours at 24h is outside of this shape, while the skin colour of our naturally tanned control group is well within the natural gamut, as expected. In general, there is a very good agreement between our baseline data (‘0h’ and ‘inn’) and the natural skin gamut. The ITAs (individual typology angles) in our control group lie between 10° (‘TS’ (tanned skin); outer forearm; Fig 5C) and 70° (‘VLS’ (very light skin); inner forearm; Fig 1B), consistent with the White (Caucasian) population data of Del Bino et al. [13], which enables us to draw robust conclusions. Comparing the ITAs of our experimental group with our own control group, the distribution of ITAs for the DHA-induced skin colours is clearly shifted to the left compared to natural tanning (Fig 4B). Crucially, the small ITAs (median 24.7°) – reflecting a decrease in lightness – are associated with an increase in the overall amount of yellowness (Fig 4A) which, in some cases, results in a skin colour outside of the natural skin colour gamut. The excessive shift to yellowness after 24 hours is consistent with the previous report by Muizzuddin et al. [10] who speculated that excessive yellowness is caused by antioxidants, or other additives contained in the DHA tanning product [21].

The maximum amount of naturally occurring yellowness is at a lightness level (*L*\*) of 50 and drops off for higher and lower skin lightness (e.g. [12]). This nonlinear co-variation in skin lightness and skin yellowness is primarily driven by the concentration of melanin in the skin which is increased by UV exposure. The corresponding DHA-induced reaction is an increase in the concentration of melanoidins, which leads to a linear co-variation of lightness and yellowness changes, instead of the curvilinear relationship characteristic of naturally occurring skin tans across ethnicities (Fig 4A).

The experimental group included six non-White individuals (one Mixed, five Asian, two of which had darker skin colour and three had lighter skin-two Chinese and one Korean-see section Participants). Their colour change followed the same trend as those of the large White sample, that is, the colour became darker and more reddish at 4h, then darker and more yellowish in 24h. While there are known mean differences in ITAs across ethnicities (e.g. Chinese, Japanese, Indian, African and White) [13], our small sample of non-whites was well within the White cluster. There was no systematic difference between female and males in our sample.

The appreciation of tanned skin seems to be contingent on various factors, such as generational, social and psychological context, and cultural background. The sunless tanning is popular in Northern Europe, Australia, North America, particularly among adolescent White female. Individuals with originally darker skin do not tend to use those products other than for clinical reasons [24]. In Asian cultures the opposite trend prevails, as evidenced by the use of skin whitening and skin lightning products [25–27]. Such social impacts of skin tanning and detailed discussion about the usage and demands are interesting topics but outside of the scope of this study. The present study focuses on the colour shift induced by the DHA. Extensive discussions about the social impacts are found in elsewhere [1, 7–9].

### The sunless tanning pathway

Colour shifts induced by the application of 7.5% concentration DHA was traced after four and 24 hours and compared with colour changes brought about by natural tanning in a control group. Comparing the orientation of the difference distributions (4h-0h), (24h-4h) and (24h-0h) reveals that the direction of the colour change is time-dependent (Fig 6). Fig 6 is plotted in Δ*a* and Δ*b*\* space, where Δ*a*\* represents the change in redness direction, Δ*b*\* represents the change in yellowish direction. After 4 hours, the skin change is more reddish than the natural tan (‘out-inn’), but between 24 and 4 hours it turns more yellowish. The resulting overall colour change (24h-0h) is well aligned with the natural tanning direction (‘out-inn’, Fig 6), but the magnitude is too great. The shift to yellowness is depicted in the skin spectral reflectances (Fig 3); after 4 hours the relative reflectance ratio (4h/0h) mimics the natural tanning ratio (out/inn), whereas a disproportionate amount of bluish light is absorbed after 24h (24h/0h) leading to a disproportionate yellowness relative to the reduction in lightness. Fig 6 shows that the shift 24h-0h is simply the vector summation of ‘4h-0h’ and ‘24h-4h’. This vectorial analysis based on PCA in the (*a*\*, *b*\*) diagram in conjunction with the spectral analysis reveals that the direction of colour shifts from the original skin to the 24 hours are very close to that of the naturally occurring tan, but that the magnitude of the shift is excessive in relationship to the lightness (not shown in Fig 6; cf. Fig 4C).

### Predicting the skin colour changes resulting from DHA-induced tanning

Application of DHA does not result in a constant colour shift for different skin samples. Assuming a constant shift in skin colour (i.e., using only the intercept as a predictor in the regression analysis) results in a median Δ*E* of 2.18 compared to a median Δ*E* of 1.87 using the regression analysis (Fig 5B). In the present study, the goodness of the prediction was represented by Δ*E* (see section Predicting the DHA-induced skin colour changes). Skin differences along the *b*\* dimension greater than 2Δ*E* can be reliably detected by colour-normal human observers when viewed side-by-side under daylight conditions; under artificial light, thresholds are higher by a factor two [28].

Very light and pale skin undergoes a more pronounced shift compared to darker skin as shown by the negative correlation in Fig 5A. This negative correlation between the observed colour shift (‘24h’) and the baseline skin colour (‘0h’) holds for all three dimensions, lightness *L*\*, redness *a*\*, and yellowness *b*\*. A stepwise regression analysis was performed using all three dimensions (*L*\*, *a*\*, *b*\*), the interactions, as well as skin moisture and pH values as predictors, but neither the interactions nor the latter two variables resulted in a substantial reduction in the variance explained in the skin colour changes (Table 2). Fig 5C shows the predicted skin colour at 24h after DHA application (black dots) using only the baseline skin colour (*L*\*, *a*\*, *b*\*) as predictors. The predicted values indicate that some variation of the resulting skin colours is captured by using the baseline skin colour as a predictor (cf. Table 2) but a substantial part of the variation is not explained by the predictors. Since we focused on the skin colour volume and ITA defined on *L*\* and *b*\*, and as shown in Fig 2A, the shift in hue varies along with time progression (more reddish after 4h, more yellowish after 24h), we chose the predictors *L*\*, *a*\*, *b*\*, rather than hue and chroma.

Of particular interest is whether we can predict the skin colours that will result in an unnatural tan, that is, a skin colour outside of the natural White skin gamut (about 20% of our sample). In Fig 7, we indicated the skin colours that lie outside the natural gamut with filled blue dots; the corresponding baseline skin samples are shown in open blue circles. The distribution of these skin colours is fairly random and cannot be predicted by our regression model or any other linear model using all colour dimensions as well as skin moisture and pH value are predictors. This reflects the individual differences in the trajectory of skin colour shift induced by DHA, which are not captured by using initial skin colour, skin moisture and pH value as predictors. Furthermore, whether a skin colour is perceived as natural is probably also determined by its consistency with other features, such as hair colour and structural facial features reflecting the individual’s ethnicity.

**Fig 7.**
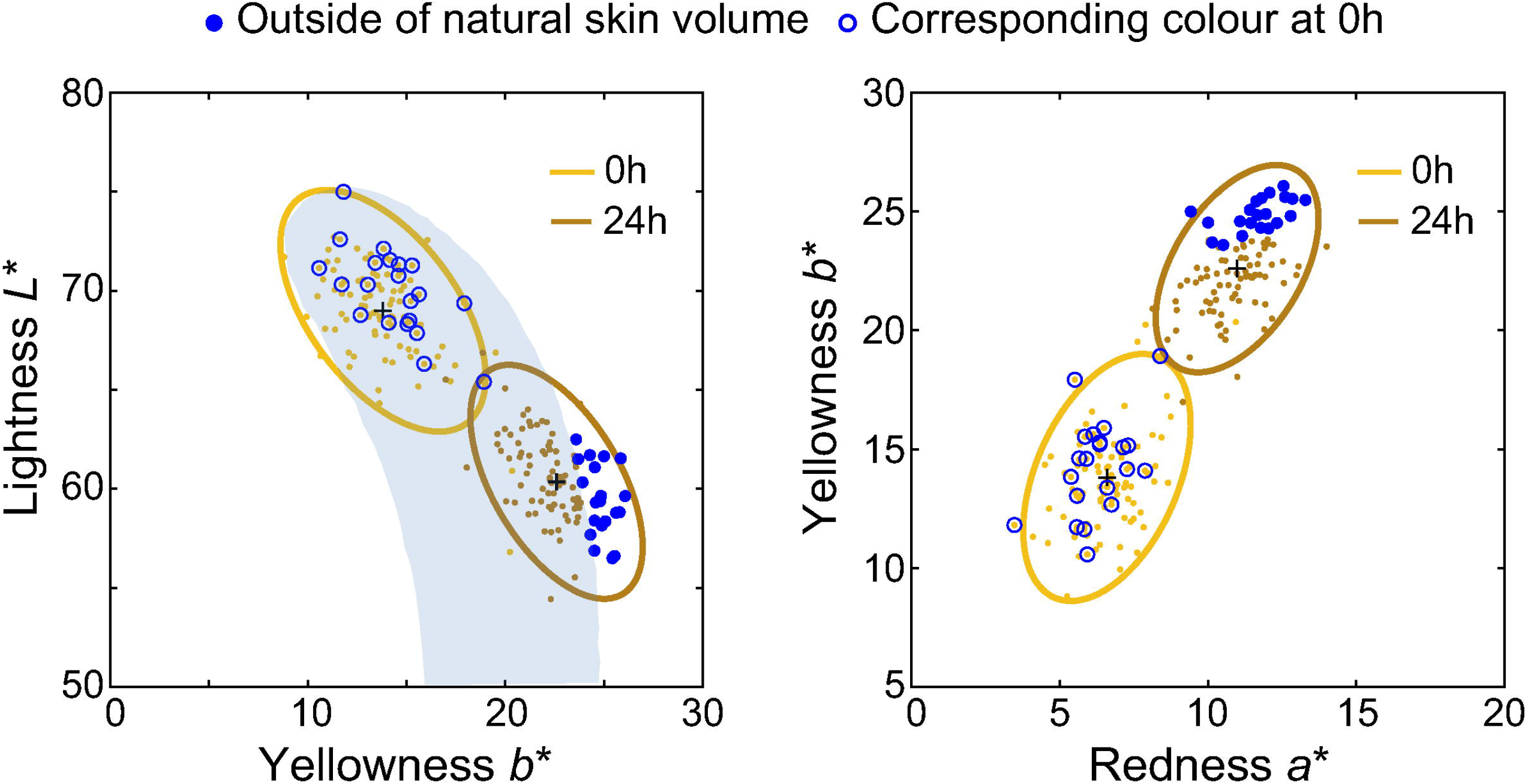
Skin tans outside of the natural skin gamut. Chromaticities of the DHA-induced skin colours outside of the ‘natural skin volume’ are shown in blue dot symbols and their original colours in blue open circle in (*L*\*, *b*\*) (left panel) and (*a*\*, *b*\*) (right panel) diagrams.

## Conclusions

Our main finding is that the direction of skin colour change induced by DHA follows a straight line in the lightness vs. yellowness diagram, leading to an excess of yellowness for a particular lightness level, while the trajectory of a natural tan is characterised by a decrease in lightness without an increase in yellowness. We conclude that the control of the colour shift magnitude is a critical factor to achieve a natural appearance, in addition to the skin hue (ITA). Secondly, we can predict with reasonable accuracy (less than 2Δ*E* median error) the overall colour shift induced by DHA from the initial skin tone. However, we cannot predict whether the skin colour of an individual will lie outside of the natural skin gamut after DHA application (about 20% of the 100 skin samples), suggesting that other individual skin differences beyond skin colour, moisture and pH value play a role in skin tanning.

## Data availability statement

Data will be made available after acceptance.

## Author Contributions

Conceptualization: KA, GM

Formal analysis: KA, GM, KX, SW

Funding acquisition: GM

Data curation: KA

Investigation: KA Methodology: GM

## Supporting information

**S1. SSV (spectral similarity value) of two spectral reflectances**

**S1 Table. Spectral similarity values between pairs of spectral reflectance**

**S2. Range of standard deviation of spectral reflectance**

**S2 Table. Range of standard deviation of spectral reflectance**

