## Supporting information for "A colorimetric comparison of sunless with natural skin tan"

**S1. SSV (spectral similarity value) of two spectral reflectances**

The similarity of spectral profiles between pairs of spectral samples can be represented by the SSV (spectral similarity value), which is the integral of Euclidian distance, corresponding to RMSE (root mean square error) and correlation between the two spectra [1-3]. Smaller SSV indicates that the two spectra are similar. SSV is zero when the two spectra are exactly the same.

Let *r*_r_ and *r*_t_ two spectra, then RMSE over wavelength (*i* ranges from 400nm to 700nm) is defined by

$RMSE= \sqrt{\frac{1}{m}\sum_{i=1}^{m} \left( r_{r,i}-r_{t,i} \right)^{2}}$,	(S1)

where *m* is the number of wavelength samples. Then, SSV is defined by

$SSV= \sqrt{{RMSE}^{2}+S^{2}}$,	(S2)

$S^{2}=1- \left[ \frac{\left\{ \frac{1}{m} \sum_{i=1}^{m} \left( r_{r, i}-\mu_{r} \right)\left( r_{t, i}-\mu_{t} \right) \right\}}{\sigma_{r} \sigma_{t}} \right]^{2}$,	(S3)

where $\mu_{r}$ and $\mu_{t}$ are means of the two spectral reflectances, and $\sigma_{r}$ and $\sigma_{t}$ are standard deviations of reflectances.

SSV are shown in S1 Table. The entries in the second column in S1 Table show the SSV calculated from individual participants: thus, SSV for corresponding participants across the conditions, then averaged over the participants. The entries in the third column were calculated from mean spectral reflectances of each condition (0h, 4h, 24h, *n* = 100; inner and outer forearm, *n* = 60).

The trends of the data are consistent with RMSE (see, section Spectral comparison between sunless and natural tans in the main text).

**S1Table. Spectral similarity values between pairs of spectral reflectance**

|  | SSV | |
| --- | --- | --- |
|  | individual spectra (SD) | mean spectra |
| 0h-4h | 0.278 (0.016) | 0.273 |
| 0h-24h | 0.321 (0.031) | 0.316 |
| inn-out | 0.275 (0.016) | 0.268 |
| 0h-inn |  | 0.255 |
| 0h-out |  | 0.274 |

**S2. Range of standard deviation of spectral reflectance**

Standard deviations (SDs) over participants across spectra (400nm-700nm) are summarised in S2 Table. The SD became smaller along with the DHA tanning developed (as time progressed from 0h to 4h and 24h). The range of SD, the minimum and maximum over wavelength in the experimental group was smaller than the control group. This could be due to the fact that the measurements of the experimental group was on the same 5 cm × 5 cm area within two days, whereas the control group on inner and outer forearms are physically different locations and the data collection of the control group were made throughout an year., and that the sample size of the experimental group was larger than the control.

**S2 Table. Range of standard deviation of spectral reflectance**

|  | SD range across 400-700nm | |
| --- | --- | --- |
|  | minimum | maximum |
| Experimental group (*n*=100) |  |  |
| 0h | 0.025 | 0.041 |
| 4h | 0.023 | 0.031 |
| 24h | 0.018 | 0.030 |
| Control group (*n*=60) |  |  |
| Inner | 0.030 | 0.089 |
| outer | 0.038 | 0.090 |
